# Mental Operations on Long-term Memory do not Require a Sustained Increase in Working Memory Engagement

**DOI:** 10.64898/2026.08.14.744869

**Authors:** İ. Efsane Algın, Eren Günseli

## Abstract

Working memory (WM) is often assumed to play a stronger role in mental operations than in pure storage. However, much of the evidence comes from tasks using novel stimuli requiring active maintenance. Everyday cognition, in contrast, often involves operating on information retrieved from long-term memory (LTM), which may not always require sustained WM storage. Moreover, prior evidence for enhanced WM involvement relies on univariate measures, which cannot separate procedural demands of operations from representational strength of operation-relevant items. Here, we used EEG to test how WM supports mental operations on LTM. First, participants studied color-position associations. Then, on each trial, a color cue prompted retrieval of its associated position, followed by a novel position. Across blocks, participants either performed a mental operation to compute the positions’ spatial midpoint or judged whether the probe matched one of the memory positions. Representations of task type and memory position were assessed using MVPA and inverted encoding models on alpha-band power, respectively. Task type was decoded throughout the trial, reflecting persistent task-set representations. In contrast, LTM position was represented in WM more strongly for integration than recognition early in the retention and operation periods, but these differences were transient. These findings challenge the view that mental operations inherently demand enhanced WM engagement: when information is available in LTM, increased WM involvement is transient, not sustained.

**Significance Statement:** Everyday activities, from planning routes to combining learned facts, require mentally operating on information stored in long-term memory. Yet how working memory, which is claimed to be central to mental operations, supports operations on existing long-term memories remains poorly understood. By dissociating item storage and task execution, this study shows that mental operations do not require persistently enhanced working memory engagement compared to recognition. Instead, working memory storage differences between tasks were transient, even though task-set representations remained distinct throughout. These findings suggest that WM can support complex mental operations on long-term memories without sustained increases beyond what is required for recognition.

## Introduction

Working memory (WM) is a system that maintains and manipulates information in an accessible state (Baddeley, 2020; Logie, 2003). This ability supports a wide range of cognitive functions and correlates with intelligence and academic success (Chota & Van der Stigchel, 2021; Unsworth et al., 2014). In particular, mental operations, such as manipulating and integrating information, have long been argued to rely critically on WM (Logie, 2003; Hyun & Luck, 2007). However, many everyday operations are performed on information that is not immediately present but must be retrieved from long-term memory (LTM) (Baddeley, 2002). For such cases, the role of WM for operations on LTM information remains unresolved.

Dual-task interference and neuroimaging findings support the role of WM in mental operations. Concurrently performing a WM task disrupts mental manipulation, suggesting shared underlying resources (Hyun & Luck, 2007; Logie, Gilhooly, & Wynn, 1994). Furthermore, mental operations are associated with increased prefrontal activity (Glahn et al., 2002; De Pisapia et al., 2007) and fronto-occipital coupling relative to mere storage (Sauseng, Klimesch, Doppelmayr, Pecherstorfer, Freunberger & Hanslmayr, 2005), regions associated with WM. These results are often interpreted as evidence that WM plays an essential role in performing mental operations.

Although many real-world operations can be performed on information available in LTM, prior research inherently necessitated sustained WM maintenance by predominantly using novel stimuli in each trial. Once information is learned, task-relevant representations can be accessed from LTM without active WM maintenance (Carlisle et al., 2011; Gunseli et al., 2014; Yılmaz et al., 2026; Liu et al., 2022). Therefore, a genuine test requires conditions where WM storage is not obligatory, but operational demands differ, allowing control-relevant processes (i.e., task-set and executive demands) to be dissociated from the storage of representations. Such a design can determine whether the greater WM involvement during mental operations reflects representational storage, task-related control, or both.

Here, we investigated this question by comparing preparation for two tasks that both require LTM retrieval but differ in whether an operation is performed on the retrieved information. Participants first learned unique color-position associations. During both tasks, a color cue prompted the retrieval of the associated position, followed by a novel position. In one condition, participants made a recognition judgment about the two positions. In the other condition, they integrated them by computing the spatial midpoint. Thus, both tasks required accurate retrieval, but only one required operating on the retrieved representation. Task conditions were blocked, allowing participants to adopt stable preparatory states associated with each task. Crucially, since the mental operation could not be performed until the second position was presented, any representational differences observed during the preceding retention interval would reflect preparatory, task-driven modulation of retrieved memory content rather than the operation itself.

To assess how and when WM is engaged when operating on information available in LTM, we drew on the high temporal precision of the EEG. We applied an inverted encoding model (IEM) to track the representational content and used multivariate analysis to decode the task type. If WM is required to store the representational content, preparing for a mental operation should lead to more robust representations of LTM-derived information compared to recognition. In contrast, if WM involvement primarily reflects procedural or control demands, then task differences should be observable only at the level of task-set representations, without necessarily modulating the strength of stimulus-specific representations. By dissociating representational and control-level contributions of WM, this approach provides a critical test of whether WM’s role in mental operations reflects the storage of content or the implementation of control, providing a principled basis for decomposing WM’s functional contributions to mental operations.

## Materials and Methods

### Participants

To estimate the required sample size, we averaged the effect sizes from studies that assessed the location selectivity of LTM-derived information using inverted encoding models applied to EEG data, included trial counts comparable to the present design, and compared CTF slopes across conditions (Sutterer et al., 2019; Foster et al., 2016; Günseli et al., 2024). Power was set to.90 and the false positive rate to.02 for an estimated effect size of d = 0.8, requiring 24 participants (G*Power 3.1.9.7; Faul et al., 2007); we therefore aimed to collect 25. If the Bayes Factor was still inconclusive at that point, we planned to keep collecting data, recomputing BFs after every five participants until BF_10_ < 0.166 or BF_10_ > 6, or until 80 participants were reached. Bayes Factors were computed using a default Cauchy prior (r = 0.707). The threshold was met at 25 participants, so data collection stopped there.

26 Sabancı University students participated in the experiment with informed consent and received course credit for their participation. The study was conducted in accordance with the Declaration of Helsinki, and ethical approval was obtained from the Sabancı University Research Ethics Committee (SUREC). One participant was excluded and replaced due to excessive ocular artifacts, resulting in a final sample of 25 participants (12 female).

### Apparatus and Stimuli

The experiment was prepared using the Psychtoolbox-3 in MATLAB (MathWorks, Natick, MA). Stimuli were presented on a reference circle (7° radius) with ten empty placeholder circles (black outlines, 1° radius) positioned equidistantly along its circumference. A fixation circle (black outlined, 0.5° radius) remained at the center of the screen throughout the experiment, changing color only when it served as a retrieval cue. The ten placeholder positions served as possible WM locations, five of which were designated as LTM positions fixed across participants. Each LTM position was associated with one of five colors: red (RGB: 255, 0, 0), purple (RGB: 18, 0, 128), cyan (RGB: 77, 190, 238), lime (RGB: 70, 255, 60), and yellow (RGB: 255, 255, 0), with color-to-position assignments randomized across participants. On each trial, the WM stimulus was a white circle (RGB: 255, 255, 255; 1° radius) presented at one of the ten placeholder positions. Probe stimuli were a black circle (1° radius) presented along the reference circle.

### Design and Procedure

The experiment consisted of three phases. In the study phase (Phase 1), participants learned five color-position associations: Each color cue was randomly paired with a specific LTM position on the reference circle (Figure 1A). Each association was presented twice for 2000 ms, jittered between 600 and 1000 ms. These associations remained fixed and were used across the whole experiment.

**Figure 1.**
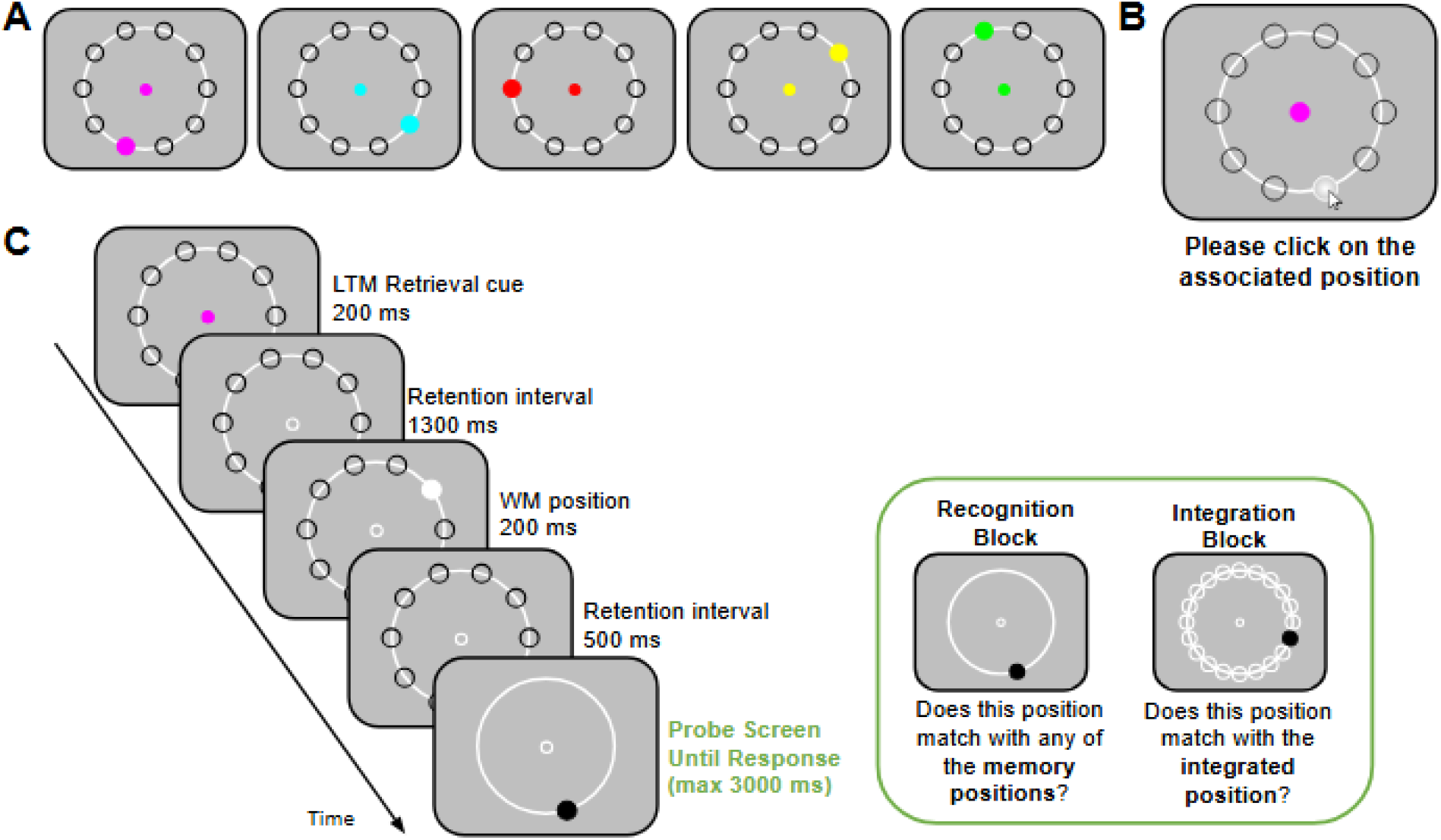
Experimental Paradigm. A. In the Study Phase, participants learned 5 color-position associations. B. In the Test Phase, these associations were tested. C. In the Experimental Phase, participants were presented with an LTM retrieval cue. Following this cue, there was a retention interval. Then, participants viewed the WM position. After a second retention interval, a probe display containing a black circle was presented. In recognition blocks, participants indicated whether the probe position matched one of the two memory positions. In integration blocks, participants indicated whether the probe position matched the integrated position, defined as the midpoint between the two memory positions along the shorter arc.

In the following test phase (Phase 2), memory for the learned associations was tested using a cued-recall procedure (Figure 1B). On each trial, a color cue appeared and remained at the center of the screen until participants clicked the associated position using the mouse. The correct color-position association was displayed after each response, regardless of accuracy. Participants advanced to Phase 3 only after achieving four consecutive correct responses for each one of the five associations.

The Experimental Phase (Phase 3) comprised 24 blocks (12 recognition, 12 integration) of 60 trials each, yielding 1440 total trials (144 per LTM position per task). Conditions alternated every four blocks, with order counterbalanced across participants. During this phase, EEG was recorded continuously, and eye position was monitored.

Each block of Phase 3 began with participants pressing the spacebar. A fixation circle remained at the center of the screen across the whole experiment, and participants were instructed to fixate on that circle (Figure 1C). After a jittered inter-trial interval (ITI; 600-1000 ms), ten empty placeholder circles positioned equidistantly along the reference circle were presented, and the fixation circle was filled and changed color to a retrieval cue (studied in Phase 1), which was presented for 200 ms. Participants were instructed to remember the associated LTM position. Following a 1300 ms retention interval, a WM display appeared for 200 ms, consisting of a white circle positioned randomly in one of the ten placeholders (WM Position), excluding both the cued LTM position and the position 180° opposite to it. After a 500 ms delay, the probe display was presented until response or for a maximum of 3000 ms.

The probe display consisted of a black circle on the reference circle. Task instructions differed by condition. In the recognition condition, participants judged whether the probe matched either of the two memory positions: LTM position, associated with the color cue, or WM position, shown via the white circle. In the integration condition, participants judged whether the probe matched the integrated position, defined as the point equidistant from the LTM and WM positions along the shorter arc of the reference circle. To ensure a shorter arc always existed, the WM position was never placed 180° from the LTM position. Responses were made using the’A’ and’D’ keys (counterbalanced across participants) to indicate’yes’ or’no’ judgments. Crucially, two tasks followed identical trial flows until the probe display. Therefore, any condition differences before probe onset can be attributed to anticipatory task preparation rather than perceptual differences.

Probe locations varied by condition. In integration blocks, probes appeared at any of 20 possible positions: the 10 original placeholders plus 10 intermediate positions between them. In recognition blocks, the probe matched one of the two memory positions (LTM or WM) on 50% of trials. The remaining 50% were mismatch trials. To make the recognition task comparably demanding to the integration task, half of the mismatch trials used a QUEST adaptive staircase procedure, which adjusted deviation magnitude (8–36°) from either the LTM or WM position. On the remaining mismatch trials, probes appeared at one of the remaining placeholder positions (>36° deviation). Feedback on response accuracy was provided after each trial.

Trials were terminated and rescheduled if participants’ gaze deviated from fixation. These terminated trials were appended to the end of the current block. At each block break, participants received accuracy feedback and reviewed the color-position associations to prevent forgetting.

At the beginning of Phase 3, participants completed a practice session with a trial structure identical to the main experiment. Practice included 40 trials (4 trials per LTM position per task). Participants were required to achieve ≥55% accuracy in both tasks to proceed. The practice session was repeated up to five times. Participants who failed to meet the criterion after the fifth attempt were planned to be excluded from the study, though all participants were able to complete the practice within five attempts (min = 55, max = 87.5, mean = 74.9).

### EEG Recording and Preprocessing

EEG was recorded from 32 Ag/AgCl electrodes (10/20 system) and mounted in an elastic cap using Brain Products actiCHamp (actiCHamp Plus, Brain Products). The signals were acquired using BrainVision Recorder (Version 1.24.0001, Brain Products GmbH). The EEG signals were amplified via an actiCHamp Plus amplifier (Brain Products GmbH, Gilching, Germany) and digitized at a sampling rate of 1000 Hz. The horizontal EOG electrodes were located 1 cm lateral to the external canthi, and the vertical EOG electrodes were located 2 cm above and below the right eye through external electrodes. The online reference electrode was located in the left mastoid (TP9), and then the data were rereferenced offline to the average of left and right mastoids (TP9 and TP10). For two participants, we placed the right HEOG onto the right mastoid due to a temporary TP10 malfunction, and rereferenced offline accordingly. The scalp montage was posterior-weighted and comprised 28 electrodes—F3, Fz, F4; FC5, FC1, FC2, FC6; C3, Cz, C4; CP5, CP1, CP2, CP6; P7, P5, P3, Pz, P4, P6, P8; PO7, PO3, PO4, PO8; O1, Oz, O2. EEG data processing was conducted using MATLAB R2022b (MathWorks, Natick, MA), along with the EEGLAB toolbox (Version 2020.1; Delorme & Makeig, 2004), the ERPLAB toolbox (Version 8.30; Lopez-Calderon & Luck, 2014), and custom scripts.

The EEG signals were filtered using a.05 - 40 Hz band-pass IIR Butterworth filter. Data were epoched to-2200 to 3800 relative to the memory-cue onset using the pop_epoch.m function from EEGLAB; the same epochs were used both for artifact rejection and subsequent time-frequency analysis. Prior to analysis, a baseline period of-200 and 0 ms relative to the memory-cue onset was subtracted from all voltage values. Artifacts related to muscle activity, slow drifts, saturation, and signal blocking were identified manually through visual inspection. For participants with usable eye tracking data (22 out of 25), ocular artifacts were monitored online and trials in which gaze deviated beyond the fixation criterion were rejected based on the saved behavioral data. For the remaining participants, ocular artifacts were inspected more carefully: trials showing a step-like saccadic deflection larger than 20 µV in the horizontal EOG (HEOG) were discarded. The artifact rejection process was performed blind to the experimental conditions and conducted prior to hypothesis testing. Trials containing artifacts were excluded, and a dataset with fewer than 80 trials per position for either condition was omitted from further analysis.

### Eye Tracking

Gaze position was monitored using a desk-mounted infrared eye-tracking system (EyeLink 1000 Plus, SR Research, Ontario, Canada), operating in remote mode. The gaze position was sampled at 500 Hz. We could track the eye position of 22 out of 26 subjects. For these participants, online feedback regarding ocular artifacts was provided throughout the experiment. Trials were aborted if the eyes deviated from fixation by 2° visual degrees throughout the sample display until the onset of the test display. This threshold was adjusted during data collection based on eye tracker data quality to optimize ocular artifact detection. For unstable gaze data, the threshold was increased to detect as many real eye movements as possible and to minimize unnecessary trial abortions due to eye tracker noise.

## Data Analysis

### Power analyses

Power analyses were performed using MATLAB’s Signal Processing toolbox (Mathworks, Natick, MA), EEGLAB toolbox (Delorme & Makeig, 2004), and FieldTrip toolbox (Oostenveld et al., 2011).The raw EEG signal was downsampled from 1000 Hz to 500 Hz and bandpass filtered to isolate frequency-specific activity at each electrode. We used a Butterworth filter as implemented by the ft_preproc_bandpassfilter.m function of the FieldTrip Toolbox. A Hilbert Transform (MATLAB Signal Processing Toolbox) was applied to the bandpass-filtered data, to extract a complex analytic signal, *z*(*t*)=*f*(*t*)+*if̃*(*t*), of the band-pass filtered EEG, *f*(*t*), where*f̃*(*t*) is the Hilbert Transform of *f*(*t*) and *i*=√−1, using the following MATLAB syntax:

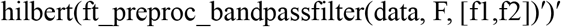

where data is the raw EEG (trials x samples), F is the sampling frequency (500 Hz), f1 and f2 are the boundaries of the frequency band to be isolated. Instantaneous total power was computed as the squared magnitude of the resulting complex signal. For the alpha frequency analysis, f1 and f2 were 8 and 12 Hz, respectively. For the theta frequency analysis, f1 and f2 were 4 and 7 Hz, respectively.

Frontal theta power was extracted from electrodes Fz and F3/4 converted to decibels (dB = 10log10(power/baseline)), using the-500 to-200 ms pre-cue interval as baseline. Bilateral alpha power was calculated similarly for occipital electrodes (Figure 3B/S2, Text S1).

**Figure 2.**
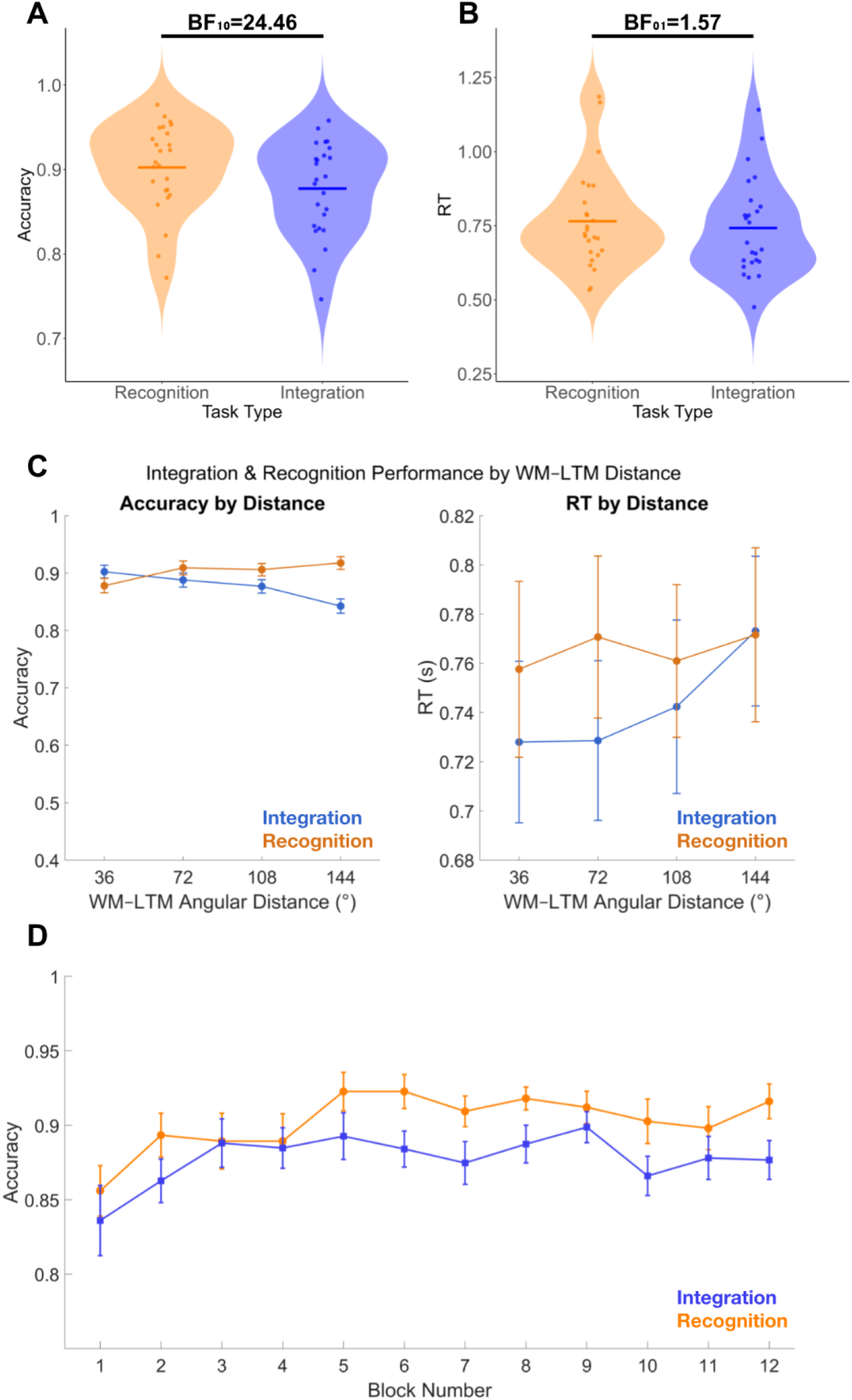
Behavioral Results for Accuracy and Reaction Times. **A.** Accuracy was higher in the Recognition than the Integration task. **B.** Reaction times did not differ between tasks. **C.** Angular distance between LTM and novel positions modulated accuracy and RT differently across tasks: in the Integration task, greater distance decreased accuracy and slowed responses, whereas in the Recognition task, accuracy increased with distance and RT was unaffected. **D.** Task accuracy across the 12 blocks of each task. Accuracy was lower for Integration (blue) than Recognition (orange) throughout the experiment, indicating that the mental operation remained more effortful. Error bars denote ±1 SEM.

**Figure 3.**
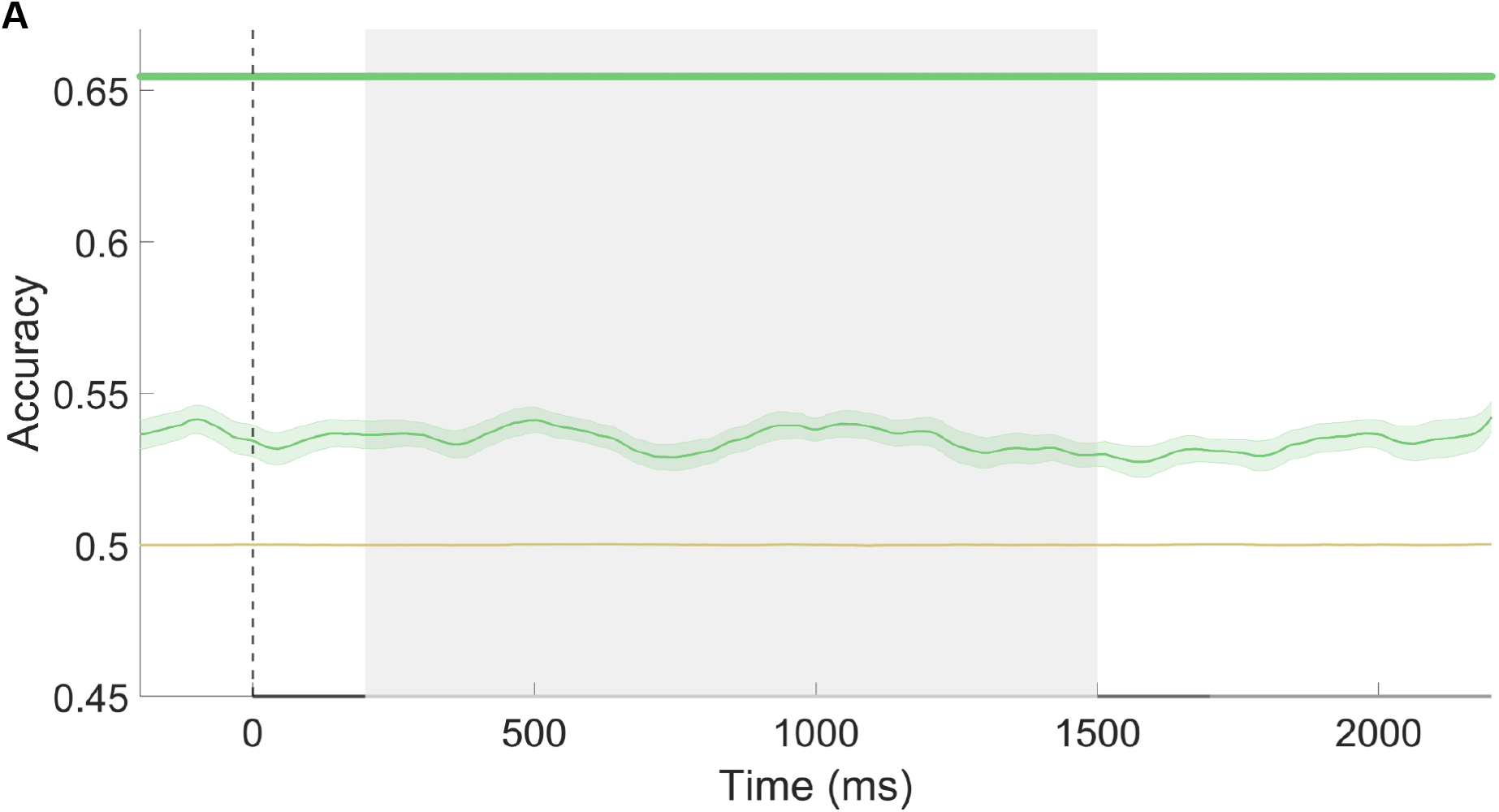
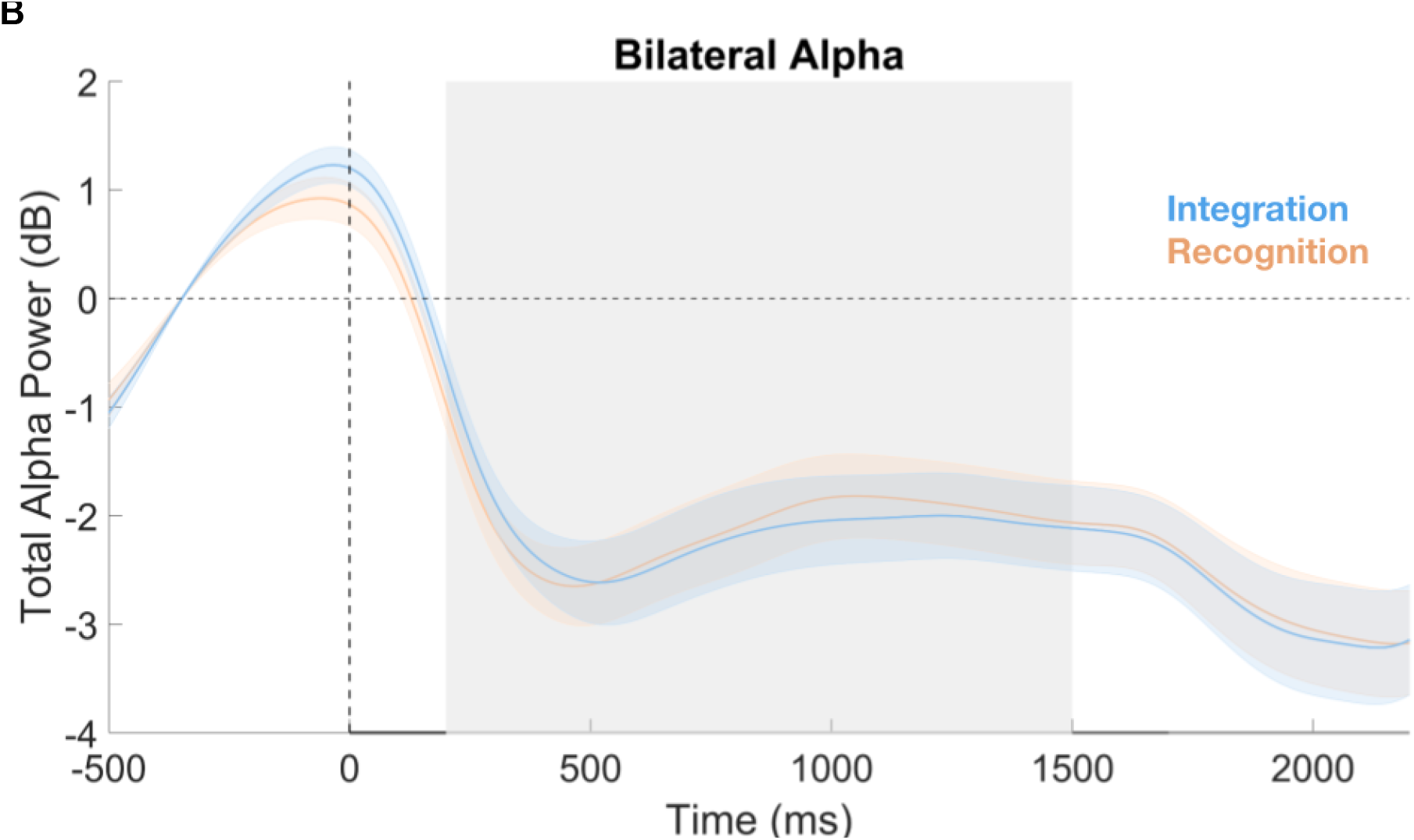
Task-type decoding from alpha power. **A.** MVPA decoding accuracy for task type (Integration vs. Recognition) based on alpha-band power patterns across the trial. Decoding remained significantly above chance throughout, including during the baseline (-200 to 0) and LTM retention (200 to 1500) periods, indicating sustained and distinct task-set representations across blocks. **B.** Total bilateral alpha power (dB) over occipital electrodes for the Integration (blue) and Recognition (orange) tasks. Alpha power did not differ between tasks at any point. Shading denotes ±1 SEM; grey marks the retention interval (200–1500 ms).

### Reconstruction of the spatial WM content

To assess the storage of spatial memory representations, we used an Inverted Encoding Model (IEM) to reconstruct spatially selective Channel Tuning Functions (CTFs) from the distribution of the total alpha-band power across posterior electrodes (C3/C4, CP1/CP2, CP5/CP6, PO3/PO4, P3/P4, P5/P6, P7/P8, PO7/PO8, O1/O2, Cz, Oz, Pz) with the assumption that each electrode reflects the weighted sum of the five spatial channels, each tuned for a different angular location (Foster et al., 2016; Günseli et al., 2024). The response profile of each spatial channel across angular locations was modeled as a half sinusoid raised to the seventh power, as:

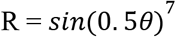

Theta is the angular location degree (ranging from 0 to 359), and R is the spatial channel response in arbitrary units. We circularly shifted the response profile for each channel such that the peak of each spatial channel response was centered over one of the five LTM positions (0°, 72°, 144°, 216°, 288°, see Figure 1).

The IEM routine was applied to each downsampled time point, proceeding in two stages (train and test). In the training stage, training data *B1* (see below) were used to estimate weights that approximate the relative contribution of the five spatial channels to the observed response measured at each electrode. Let *B1* (*m* electrodes × *n1* observations) be the power at each electrode for each measurement in the training set, *C1* (*k* channels × *n1* observations) be the predicted response of each spatial channel (determined by the basis functions) for each measurement, and W (*m* electrodes × *k* channels) be a weight matrix that characterizes a linear mapping from “channel space” to “electrode space”. The relationship between *B1*, *C1*, and *W* can be described by a general linear model of the form:

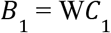

The weight matrix was obtained via least-squares estimation as follows:

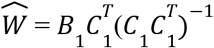

In the test stage, with the weights in hand, we inverted the model to transform the observed test data *B2* (*m* electrodes × *n*_2_ observations) into estimated channel responses, *C2* (*k* channels × *n*_2_ observations):

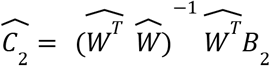

We used a “leave-one-out” cross-validation routine such that two blocks of estimated power values (see

Power Analysis) served as *B*_1_ and were used to estimate Ŵ, and the remaining block served as *B*_2_ and was used to estimate *n*_2_, ensuring that the training and test data were always independent. This process was repeated until each of the three blocks was held out as the test set, and the resulting CTFs were averaged across each test block. This IEM routine was applied separately for each subject, and statistical analyses were performed on the reconstructed CTFs.

Finally, because we equated the number of trials across position bins within blocks, a random subset of trials was not included in any block. Thus, we randomly generated 100 block assignments, each resulting in a *p*∗*b × m × s* power matrix. The IEM routine was applied to the matrices of power values for each block assignment, and their outputs (i.e., channel response functions) were averaged. This iterative approach better utilizes the complete dataset for each subject and reduces noise in the resulting CTFs by minimizing the influence of idiosyncrasies in estimates of power specific to any given assignment of trials to blocks.

To quantify the location selectivity of the CTFs, we estimated the CTF slope (i.e., the “peakedness” of the channel response at channel offset zero relative to farther channel offsets). The CTF slope was estimated using linear regression by collapsing the channel responses across channels that were equidistant.

The IEM was applied separately to LTM and WM positions. For LTM positions, five spatial channels were used, each centered on one of the five learned positions. For WM positions, to obtain sufficient trial counts for stable IEM estimation, the 10 possible positions were grouped into five bins of two adjacent positions, as each individual position was repeated only 60-70 times. To ensure that tuning estimates were not biased by arbitrary grouping choices, this procedure was repeated with two offset grouping schemes (e.g., positions 1-2, 3-4, 5-6, 7-8, 9-10 in the first iteration, and positions 2-3, 4-5, 6-7, 8-9, 10-1 in the second), and CTF slopes were averaged across iterations to obtain the final CTF slope estimates. For integrated/midpoint positions, we calculated the shortest angular distance (shorter arc) between each LTM position and WM position, and computed the midpoint of this distance. To obtain sufficient trial counts, we followed a similar approach to WM positions but grouped four adjacent positions together because there were 20 possible midpoint positions. As before four offset grouping schemes were iterated over to ensure unbiased estimates. We then computed CTF slopes for each grouping scheme and averaged across iterations to obtain the final CTF slope estimates. For LTM, WM, and integrated positions, the IEM routine was applied separately for Recognition and Integration tasks.

### Cross-Task Inverted Encoding Model

To test whether spatial representations generalize across task contexts, we performed a cross-task IEM analysis. We trained the encoding model on one task condition and tested it on the other task condition, examining whether spatial channel tuning functions could be reconstructed across different task demands.

For each iteration, two blocks from one task condition (e.g., Integration) were used to train the weight matrix, and one block from the other task condition (e.g., Recognition) was used to reconstruct channel responses. This ensured that training and test data came from different task contexts. This procedure was performed in both training and testing directions.

As in the within-task analysis, we equated the number of trials across position bins within blocks and randomly generated 100 block assignments to match the number of testing iterations in the within-task IEM analysis (100 iterations × 3 cross-validation folds). The IEM routine was applied for each block assignment, and the resulting channel response profiles were averaged. CTF slopes were computed for each decoding direction and averaged to obtain the final cross-task CTF slope estimates.

### Multivariate Pattern Analysis

We conducted a decoding analysis to assess whether the representations of task sets differed (operation and recognition) using the CoSMoMVPA toolbox (Oosterhof et al., 2016) for MATLAB. The analysis was conducted separately for each participant, with a linear discriminant analysis (LDA) classifier trained and tested on the pattern of total alpha power across all available EEG electrodes, averaged within non-overlapping 10 ms temporal windows. Specifically, we adopted a 10-fold cross-validation scheme in which the LDA classifiers were trained on 9 folds and tested on the held-out fold with each fold serving as the test set once. This procedure was repeated across 100 iterations of random fold assignments.The classification accuracy was calculated as the proportion of correct predictions averaged across all folds and iterations.

## Statistical Comparisons

We adopted two main approaches for statistical comparisons of EEG data: first, whether location selectivity was present (average CTF vs. permuted null); second, whether location selectivity differed across conditions (integration vs. recognition). For both comparisons, we averaged CTF slopes over the LTM retention interval (200–1500 ms) and computed Bayesian paired-samples t-tests. The same approach was applied to WM positions, with CTF slopes averaged over the second retention interval (1700–2200 ms).

We performed Bayesian and frequentist paired-samples t-tests, and repeated measures ANOVAs for all comparisons using JASP (JASP Team, 2019) using the default priors, with effects evaluated across the model space (“across all models”). When evidence favored the alternative hypothesis, we reported BF₁₀; when it favored the null, we reported BF₀₁ = 1/BF₁₀. Bayes factors were interpreted using standard thresholds: 1–3 anecdotal, 3–10 moderate, 10–30 strong, 30–100 very strong, >100 extreme (Lee & Wagenmakers, 2014; see also Schönbrodt & Wagenmakers, 2018; Dienes, 2021).

To obtain permuted null CTF slopes, assess whether CTF slopes were statistically different from chance, we first created a null distribution by randomizing the location labels within each block so that the labels were random with respect to the observed EEG signal in each electrode. This randomization procedure was repeated 1,000 times to obtain permuted null CTF slopes for each participant.

To examine the temporal dynamics of potential task differences, we performed a cluster-based permutation test (Maris & Oostenveld, 2007) comparing CTF slopes between integration and recognition at each time point, implemented in FieldTrip (Oostenveld et al., 2011) with a Monte Carlo randomization procedure. This non-parametric approach was employed as the mean CTF slope may not be normally distributed under the null hypothesis. We ran two separate tests. The first spanned cue onset through the LTM retention interval (0 to 1500 ms), the second from WM onset to the probe (1500 to 2200 ms), when the mental operation could take place in the integration task. Where a significant cluster was identified, we confirmed the effect by averaging CTF slopes within the identified time window and performing a Bayesian paired-samples t-test. To test whether the tasks converged after this interval, we averaged CTF slopes over the remaining retention period and performed the same test.

For all comparisons, temporally adjacent data points with a p-value smaller than.05 were clustered together. Condition labels were then randomly shuffled for 10,000 iterations. At each iteration, independent-samples t-tests were performed on shuffled data. A cluster-level statistic was calculated by taking the sum of the t-values within each cluster, separately for positive and negative clusters. The p-value for each cluster was calculated as the proportion of permutations in which the maximum cluster-level statistic under random shuffling exceeded that of the observed cluster. A cluster was considered significant if the calculated p-value was smaller than.05. This approach corrects for multiple comparisons by controlling the family-wise error rate across temporal comparisons (Maris & Oostenveld, 2007).

Cluster-based permutation tests comparing CTF slopes against permuted null values separately for each task condition are reported in Table 2 (Figure 4, 5 & 6).

**Figure 4.**
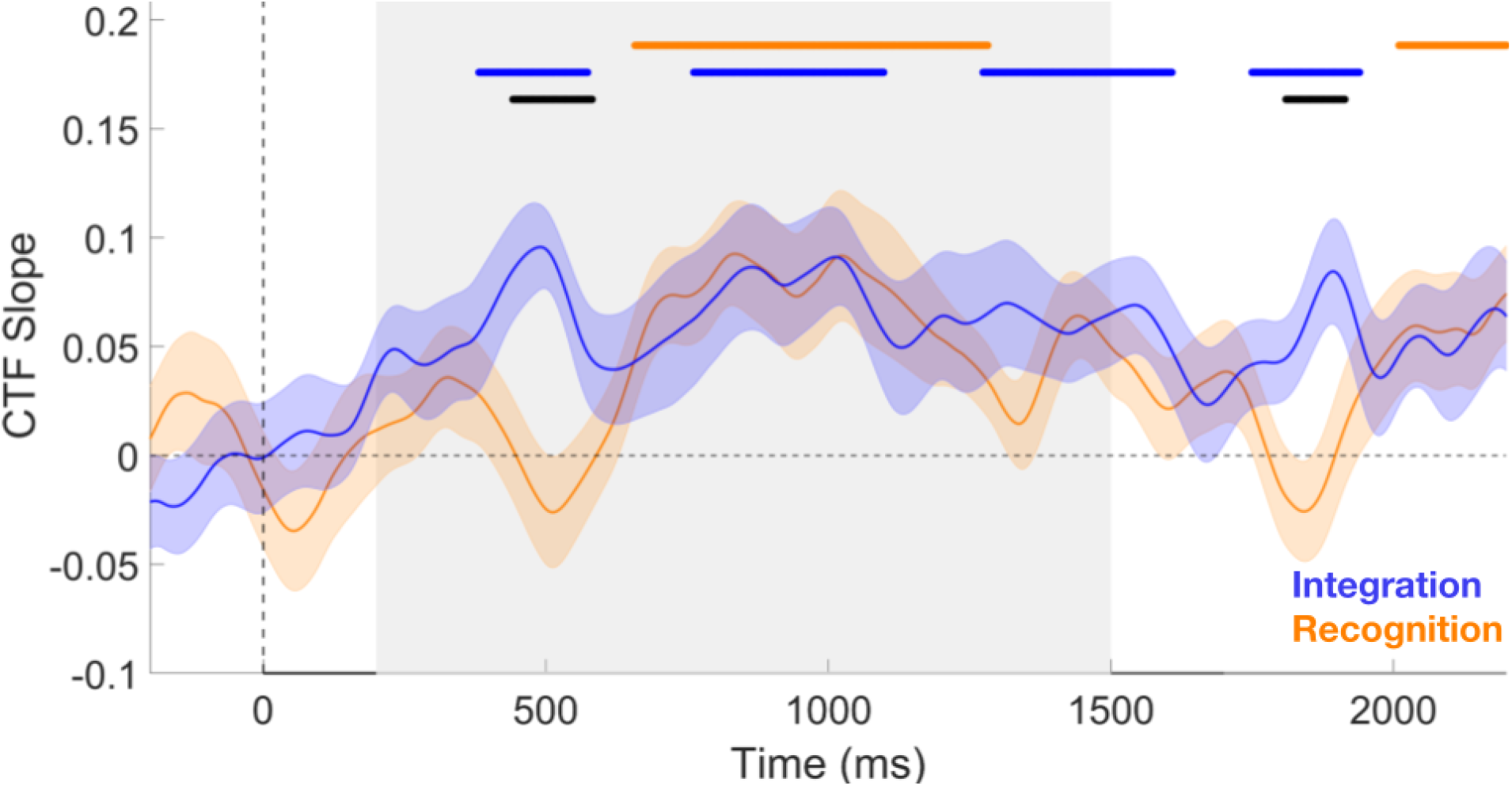
LTM CTF slopes by task. The location selectivity for LTM positions is shown separately for the Integration task and the Recognition task. The grey-shaded region marks the retention interval from 200 to 1500 ms. The black lines above represent the time windows 442 to 582 ms and 1810 to 1914 ms, respectively, identified through cluster-based permutation testing as showing a significant difference across the conditions. For the recognition task, two significant positive clusters emerged: the first from 658 to 1282 ms and the second from 2010 to 2486 ms (ps <.05; two-tailed). For the integration task, Four significant positive clusters emerged: the first from 382 to 574 ms, the second from 762 to 1098 ms, the third from 1274 to 1608 ms and the fourth from 1750 to 1940 ms (ps <.05; two-tailed).

**Figure 5.**
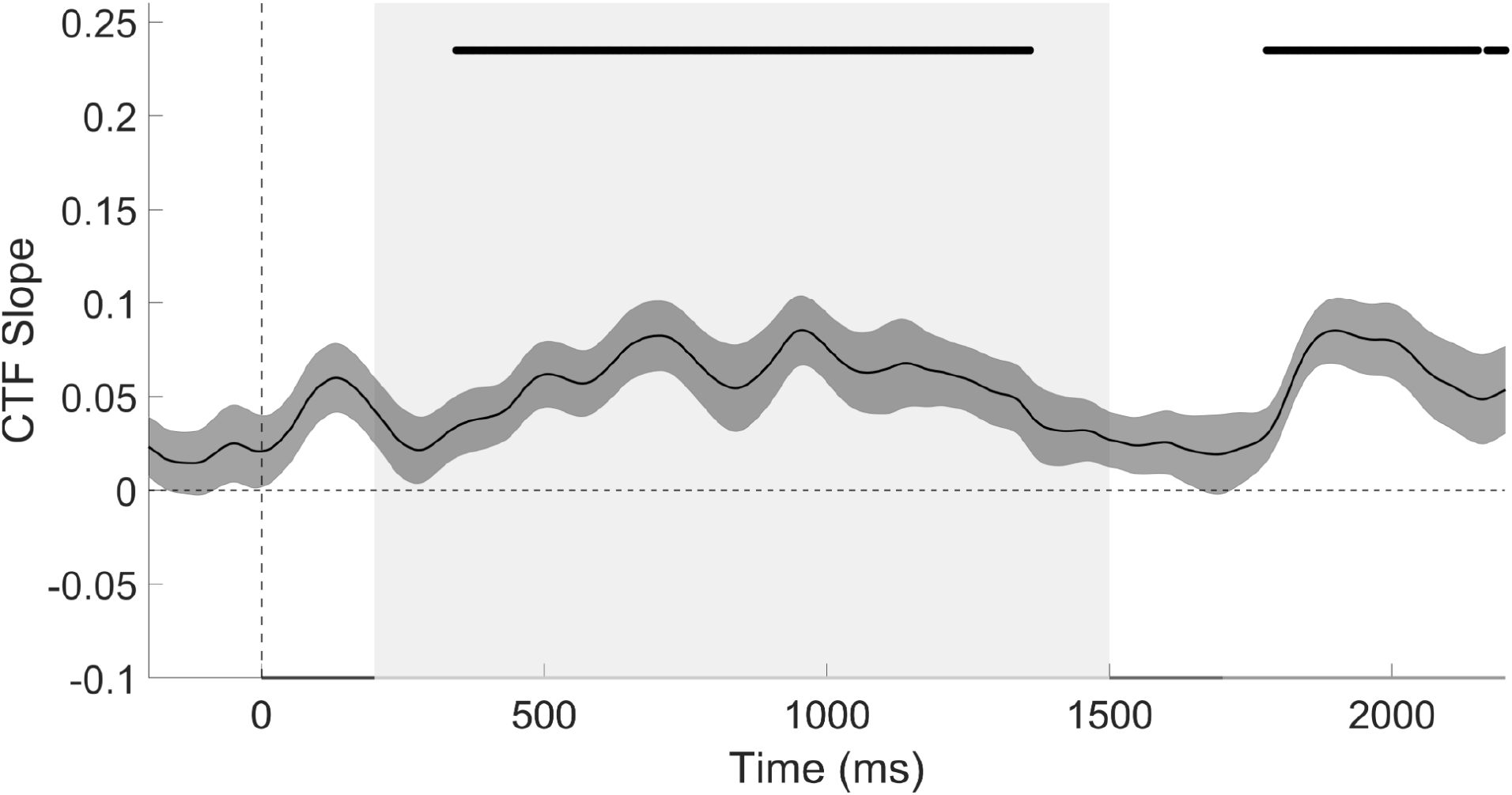
Cross-Task CTF Slopes. Cross-task CTF slopes for LTM positions, averaged across both directions (trained on one task, tested on the other). CTF slopes were above chance across the retention interval (grey shaded, 200 to 1500 ms), indicating that LTM position representations generalized across the Integration and Recognition tasks. To complement the retention-interval average analysis reported in the main text, we used cluster-based permutation testing to examine the time course of cross-task CTF slopes against permuted (null) CTF slopes. Three significant positive clusters emerged: the first from 344 to 1360 ms, the second from 1778 to 2152 ms, and the third from 2168 to 2376 ms (ps <.05; two-tailed).

## Results

### Behavioral

Behavioral results are provided in Figure 2 and Table S1. The accuracy of participants was significantly higher for the Recognition task (*M* = 90.2%, *SD* =5.2%) than the Integration task (*M* = 87.7%, *SD* = 5.5%), *t*(24) = 3.585, *p =*.001, BF₁₀ = 24.46. However, the reaction times did not differ between the Recognition (*M* = 765 ms, *SD* = 166 ms) and Integration (*M* = 742 ms, *SD* = 161 ms) tasks, *t*(24) = 1.588, *p =*.125, BF₀₁ = 1.57. These results suggest that the Integration task was harder despite using an adaptive staircasing procedure to match the accuracy across tasks.

As an exploratory analysis of task demands, we inspected accuracy and RT as a function of the angular distance between the LTM and novel positions across trials (Figure 2C). We conducted frequentist and Bayesian repeated-measures ANOVAs with Task Type (integration, recognition) and Angular Distance (36°, 72°, 108°, 144°) as within-subject factors. For Accuracy, Mauchly’s test indicated a violation of sphericity for the Task Type × Angular Distance interaction, χ²(5) = 12.89, p =.02, and degrees of freedom were corrected using the Huynh-Feldt estimate (ε = 0.78). Results revealed extreme evidence for main effects of Task Type, BF₁₀=4.58e12, *F*(1, 24) = 12.825, *p* =.002, η*_p_*^2^ = 0.35, and Angular Distance, BF₁₀ = 3.2e11, *F*(2.65, 63.575) = 3.759, *p* =.019, η*_p_*^2^ = 0.14 as well as their interaction, BF₁₀ =1.39e12, *F*(2.35, 56.34) = 25.053, *p* <.001, η*_p_*^2^ = 0.51. Follow-up analyses revealed that Angular Distance yielded a linear effect on accuracy in both tasks, *t*(24) =-9.877, *p* <.001, *d* =-0.715 for integration and *t*(24) = 4.133, *p* <.001, *d =* 0.444 for recognition, but in opposing directions: accuracy decreased with increasing distance in the integration task, whereas it increased in the recognition task. RT results partially mirrored the pattern of accuracies revealing a significant main effect of Angular Distance, BF₁₀ =28.05, *F*(3, 72) = 6.95, *p* <.001, η*_p_*^2^ = 0.225 and its interaction with TaskType, BF₁₀ =7.75, *F*(3, 72) = 3.331, *p* =.026, η*_p_*^2^ = 0.122, but not Task Type, *F*(1, 24) = 3.33, *p* =.137. In the integration task, RTs linearly slowed with increasing angular distance, *t*(24) = 4.53, *p* <.001, *d* = 0.203, whereas angular distance had no significant effect on recognition RTs, *F*(3, 72) = 1.06, *p* =.372. These results indicate that the angular distance between LTM and novel positions modulated behavioral performance differently across tasks. For integration, greater distances likely increased the demands of computing the midpoint between the two positions, leading to lower accuracy and slower responses, while recognition performance benefited from increased spatial discriminability between the two positions (see Table 1 for pairwise comparisons).

**Table 1.** Pairwise comparisons for angular difference across positions for Accuracy and Reaction Times.

| Accuracy |  |  |  |  |  |  |  |  |
| --- | --- | --- | --- | --- | --- | --- | --- | --- |
| Comparison | Recognition |  |  |  | Integration |  |  |  |
| | $BF_{10}$ | $t(24)$ | $p_{holm}$ | Cohen's $d$ | $BF_{10}$ | $t(24)$ | $p_{holm}$ | Cohen's $d$ |
| 36° vs 72° | 4.73 | -2.79 | .152 | -0.531 | 1.16 | 1.999 | .57 | 0.246 |
| 36° vs 108° | 3.52 | -2.64 | .202 | -0.475 | 16.79 | 3.41 | .044 | 0.434 |
| 36° vs 144° | 180.72 | -4.48 | .004 | -0.672 | 7.82e6 | 9.45 | <.001 | 1.02 |
| 72° vs 108° | 0.25 | 0.61 | 1 | 0.056 | 0.54 | 1.46 | 1 | 0.187 |
| 72° vs 144° | 0.34 | -1.03 | 1 | -0.141 | 17.62e3 | 6.5 | <.001 | 0.770 |
| 108° vs 144° | 1.05 | -1.94 | .581 | -0.197 | 91.9 | 4.18 | .007 | 0.582 |
| Reaction Times |  |  |  |  |  |  |  |  |
|  | Recognition |  |  |  | Integration |  |  |  |
| | $BF_{10}$ | $t(24)$ | $p_{holm}$ | Cohen's $d$ | $BF_{10}$ | $t(24)$ | $p_{holm}$ | Cohen's $d$ |
| 36° vs 72° | 0.39 | -1.182 | 1 | -0.08 | 0.21 | -0.056 | 1 | -0.004 |
| 36° vs 108° | 0.23 | -0.427 | 1 | -0.02 | 0.51 | -1.417 | 1 | -0.09 |
| 36° vs 144° | 0.91 | -1.841 | 1 | -0.08 | 126.3 | -4.323 | .007 | -0.27 |
| 72° vs 108° | 0.31 | 0.914 | 1 | 0.06 | 0.51 | -1.418 | 1 | -0.08 |
| 72° vs 144° | 0.21 | -0.077 | 1 | -0.005 | 111.07 | -4.266 | .007 | -0.27 |
| 108° vs 144° | 0.44 | -1.283 | 1 | -0.064 | 9.52 | -3.14 | .115 | -0.19 |

**Table 2.** Paired samples t-tests for CTF Slopes.

| **LTM positions averaged over the first retention interval (200 to 1500 ms).** |  |  |  |  |  |
| --- | --- | --- | --- | --- | --- |
| **Condition** | **BF<sub>10</sub>** | **95% Credible Interval** | ***t*(24)** | ***p*** | **Cohen's *d*** |
| Recognition vs Null | 121.8 | [0.394 1.316] | 4.744 | <.001* | 0.861 |
| Integration vs Null | 1698.39 | [0.587 1.583] | 5.460 | <.001* | 1.092 |
| **LTM positions averaged over the second retention interval (1700 to 2200 ms).** |  |  |  |  |  |
| **Condition** | **BF<sub>10</sub>** | **95% Credible Interval** | ***t*(24)** | ***p*** | **Cohen's *d*** |
| Recognition vs Null | 0.926 | [-0.038 0.773] | 1.855 | .076 | 0.371 |
| Integration vs Null | 7.88 | [0.176 1.032] | 3.048 | .006* | 0.610 |
| Integration vs Recognition | 0.382 | [-0.625 0.169] | 1.153 | .26 | 0.231 |
| **WM positions averaged over the second retention interval (1700 to 2200 ms).** |  |  |  |  |  |
| **Condition** | **BF<sub>10</sub>** | **95% Credible Interval** | ***t*(24)** | ***p*** | **Cohen's *d*** |
| Recognition vs Null | 186495 | [0.931 2.09] | 7.588 | <.001* | 1.518 |
| Integration vs Null | 304341 | [0.968 2.146] | 7.822 | <.001* | 1.564 |
| Condition Average vs Null | 643732 | [1.025 2.234] | 8.185 | <.001* | 1.637 |

## EEG

### Multivariate Pattern Analysis

#### Sustained representation of task rules across the trial

MVPA results are shown in Figure 3A. MVPA revealed sustained above-chance decoding of task types (Integration vs. Recognition) based on alpha-band power patterns throughout the trial (cluster p <.05). Given the blocked task design, this sustained discriminability was expected. Importantly, the two tasks followed identical sequences until the onset of the second memory position, after which participants could engage in mental integration or maintenance. Therefore, to isolate the preparatory activity from task-specific processing, we focused on the retention period after LTM-cue onset (0 to 1500 ms), preceding the presentation of WM position. Additionally, we examined the baseline period (-200 to 0 ms), to avoid contamination from the differences in stimulus-evoked activity. There was extreme evidence for above chance decoding accuracy during the baseline period, BF₁₀ = 8.9 5, *t*(24) = 8.345, *p* <.001, and retention interval, BF₁₀ = 6.17 7, *t*(24) = 10.577, *p* <.001. These results suggest that participants stored distinct task sets across blocks. Notably, univariate alpha power did not differ between tasks (Figure 3B, BF₀₁ = 4.23, *t*(24) = 0.502, *p* =.620), indicating that task discrimination relied on distributed spatial patterns rather than overall amplitude differences.

#### Inverted Encoding Model

LTM positions were represented more robustly during the integration task for the early retention interval

IEM results are provided in Figure 4 and Table 2. We first assessed whether position representations could be reconstructed from alpha-band activity during the retention interval. CTF slopes averaged across the entire retention interval provided extreme evidence for successful location selectivity, *t*(24) = 5.6, *p* <.001, BF₁₀ = 2349.22, confirming that IEMs applied to EEG data can reconstruct spatial representations retrieved from LTM (Sutterer et al., 2019).

To compare preparatory location selectivity across tasks, we first compared CTF slopes averaged across the entire retention interval, which did not differ between tasks, BF₁₀ = 0.781, *t*(24) = 1.739, *p* =.095. We then conducted a cluster-based permutation test to examine potential temporal differences, which revealed that location selectivity was higher for the integration task compared to the recognition task during the early retention interval (442 to 582 ms from cue onset). To further test this observation, we compared averaged CTF slopes during this time interval. There was strong evidence for larger CTF slopes in the Integration blocks, BF₁₀ = 16.18, *t*(24) = 3.393, *p* =.002. These findings suggest that the LTM positions were more robustly represented early in the retention interval for the Integration task.

### LTM position representations converged to the same levels for both tasks

To avoid contamination from the early task-specific difference and the second position onset, we extracted and averaged CTF slopes for each task separately within the time window from 582 to 1500 ms. There was moderate evidence for equal CTF across task types, BF₀₁ = 4.53, *t*(24) = 0.319, *p* =.752. This finding suggests that WM engagement to store retrieved LTM memory positions was overall comparable across tasks.

#### LTM position representations were stronger again for the integration task during the operation period

We examined whether CTF slopes for LTM positions differed between tasks during the period when the mental operation was expected to take place (1500 to 2200 ms from cue onset). A cluster-based permutation test revealed a significant cluster in which CTF slopes were higher for the integration task compared to the recognition task (1810 to 1914 ms from cue onset). Averaged CTF slopes within this window provided strong evidence for larger CTF slopes in the integration blocks, BF₁₀ = 6.16, *t*(24) = 2.926, *p* =.007. This suggests that LTM representations were more robust during the mental operation itself. Importantly, the brevity of the significant clusters reflects the transient nature of the effect rather than a lack of statistical control, since cluster-based permutation already corrects for multiple comparisons across the full time course.

#### LTM position representations were shared across tasks

To examine whether spatial representations transferred across task contexts, we performed a cross-task IEM analysis (Figure 5). CTF slopes were averaged across the retention interval (200-1500 ms) and compared against permuted CTF slopes using a Bayesian paired-samples t-test. Results revealed extreme evidence for location selectivity, BF₁₀ = 1356, *t*(24) = 5.361, p <.001, indicating that spatial representations were retrieved in a task-invariant format that generalized across Integration and Recognition conditions.

#### Location selectivity for WM positions did not differ across conditions except a brief late cluster

Although our main objective was to assess the representational strength for LTM positions, we conducted a complementary analysis to measure WM representations during the second retention interval. We averaged CTF slopes for WM positions during this interval (1700 to 2200 ms). A paired-samples t-test provided anecdotal evidence in favor of equal CTF slopes, BF₀₁ = 2.69, *t*(24) = 1.127, *p* =.271, suggesting that WM positions were represented with comparable location selectivity across the Integration and Recognition tasks. However, this result should be interpreted with caution because participants could already start performing the mental operation in Integration blocks, potentially replacing the original WM position with the computed midpoint. Consistent with this possibility, cluster-based permutation test across conditions revealed that the location selectivity for WM positions was briefly lower in the Integration task than in the Recognition task between 2058 and 2160 ms, BF₁₀ = 4.3, *t*(24) = 2.742, *p* =.011 (Figure 6).

**Figure 6.**
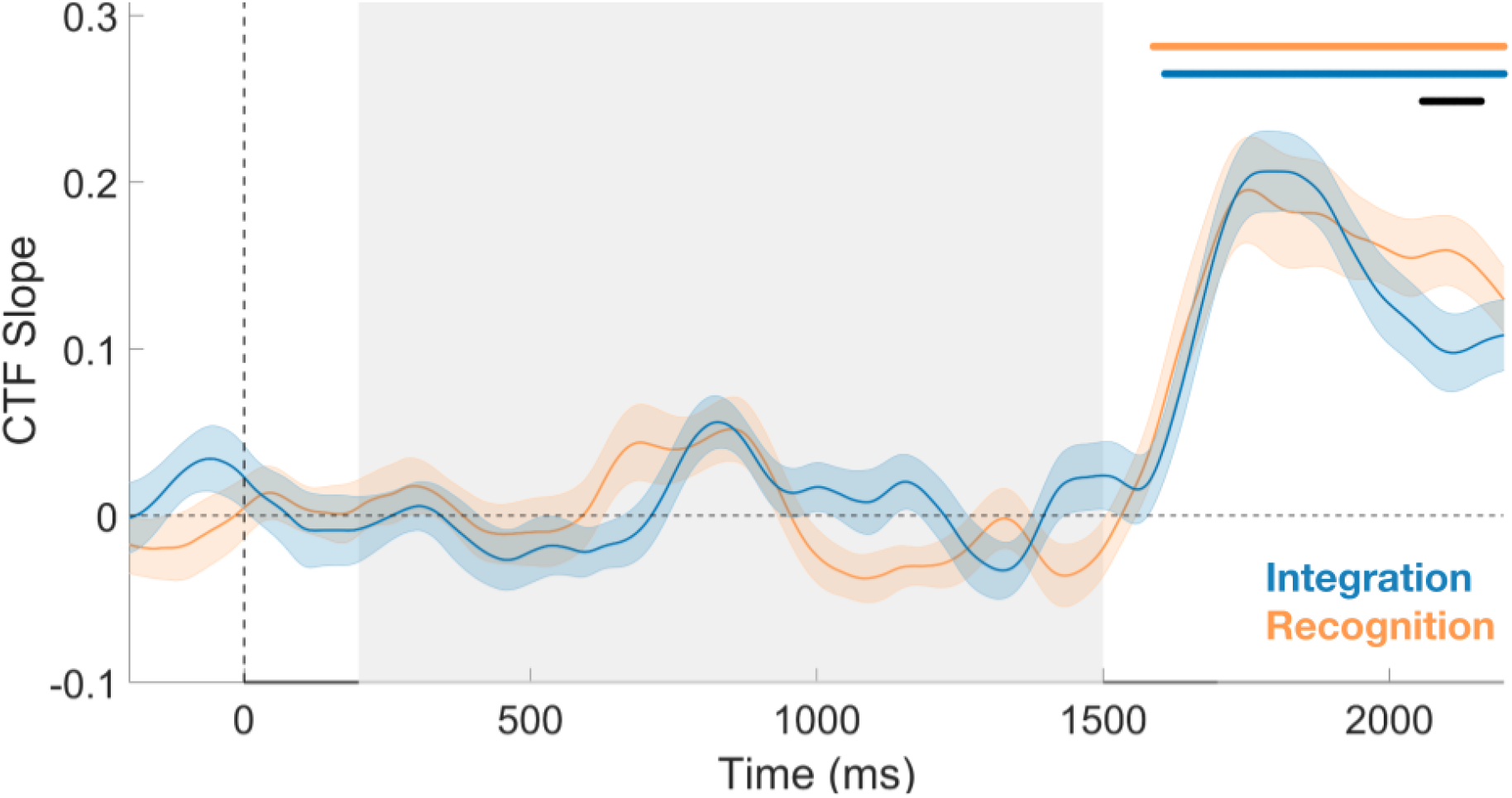
WM CTF slopes: within-and between-task comparisons (CBPT) CTF slopes for WM positions in the Integration and Recognition tasks across the trial. Slopes rose after the WM position appeared (1500 ms) and did not differ between tasks during the second retention interval (1700 to 2200 ms). We used cluster-based permutation testing to examine the time course of WM CTF slopes against permuted (null) CTF slopes within each task condition. For the recognition task, one significant positive cluster emerged from 1588 to 2200 ms (ps <.05; two-tailed). For the integration task, one significant positive cluster emerged from 1608 to 2200 ms (ps <.05; two-tailed). The black bar marks a brief late cluster (2058 to 2160 ms) in which slopes were lower for Integration than Recognition.

#### Frontal Theta power did not differ across conditions

To assess potential differences in attentional engagement and procedural demands across tasks, we examined frontal theta power (4–7 Hz) (Itthipuripat et al., 2013). A cluster-based permutation test revealed no significant clusters throughout the trial, indicating that frontal theta power did not differ across the integration and recognition tasks. We further compared averaged theta power during the second retention interval (1700–2200 ms), when the mental operation was expected to take place in the integration task. There was moderate evidence towards the null hypothesis, BF₀₁ = 4, *t*(24) = 0.614, *p* =.545 (Figure 7).

**Figure 7.**
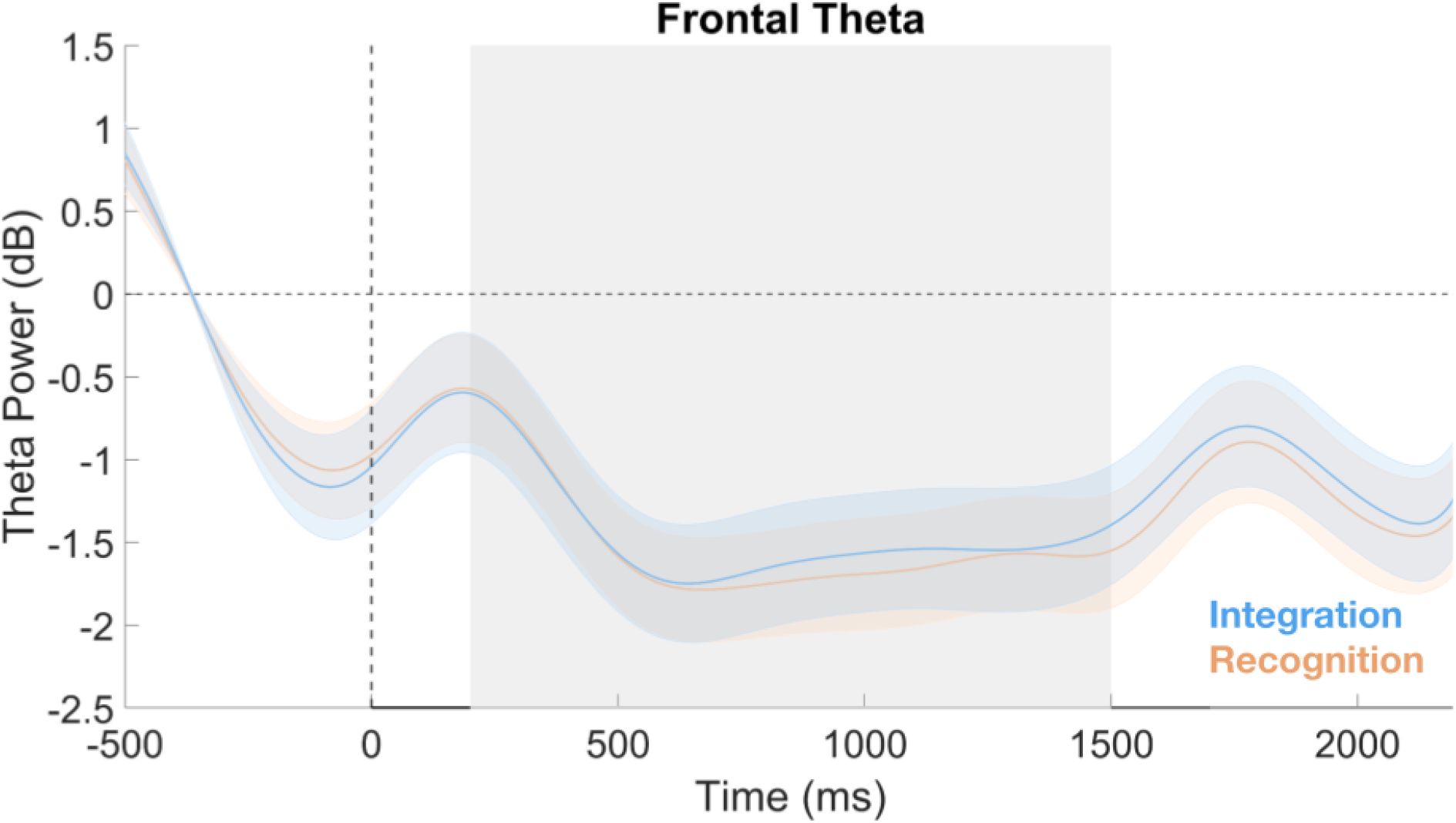
Frontal theta power by task. Frontal theta power (dB; 4–7 Hz; F3/4/z) for the Integration and Recognition tasks across the trial. Theta power did not differ between tasks at any point, and the Bayesian analysis favored no difference during the operation period (1700 to 2200 ms).

## Discussion

In the current study, we investigated the role of WM for performing mental operations on information retrieved from LTM. Participants prepared for integration (mental operation) or recognition tasks involving spatial positions. Multivariate pattern analysis confirmed that the two tasks elicited distinguishable preparatory neural patterns. To index WM storage, we applied an IEM to alpha-band EEG activity and quantified location selectivity using CTF slopes. WM engagement levels to represent retrieved LTM representations were comparable across tasks, except for an initially more robust engagement when preparing for integration than recognition, a difference that disappeared toward the end of the retention interval. Larger location selectivity for integration re-emerged for a short period when the operation was expected to take place. These results demonstrate that during preparation, and even during the operation itself, LTM information stored for operation and recognition were represented with mostly comparable WM involvement except for brief periods. Our findings suggest that mental operations may not necessarily demand stronger sustained WM involvement than mere recognition when operating on information retrieved from LTM.

Higher CTF slopes early in the retention and during operation could reflect representations that are either stronger (higher gain) or more precise (finer tuning). We argue that this difference primarily reflects gain rather than precision, for several reasons. First, while the probe position for the Integration task was selected among fixed placeholders, for the Recognition task, the probe was slightly deviated (8–36°) from either LTM or WM positions on 25% of trials, arguably requiring finer spatial discrimination. Given that probes requiring finer discrimination can enhance precision (Machizawa et al., 2012; Master et al., 2024; Allon et al., 2014), if CTF slopes mainly reflected memory precision, both LTM and WM slopes should have been higher for recognition vs integration blocks, which was not the case. Furthermore, the location placeholders remained visible throughout the trial, allowing equal precision for LTM representations across both tasks once retrieved. Taken together, CTF slope differences are more likely to reflect differences in memory reactivation strength, with WM briefly recruited more for storing the LTM contents in integration than recognition blocks.

The transient representational difference may arise because integration imposes greater computational demands than recognition and encourages participants to proactively establish item-task associations before the operation can begin. These demands could give rise to the observed neural differences through at least two complementary mechanisms. First, participants may have adopted a flexible strategy that briefly relied on latent representational states, such as LTM or activity-silent short-term storage (Ataseven et al., 2026; Günseli, in prep.; Risko & Gilbert, 2016; Yılmaz et al., 2026; Wolff et al., 2017). Paralleling cognitive offloading in the external domain, such “internal offloading” could help minimize cognitive and metabolic costs for storing items that will be used in the less demanding recognition task. Second, WM engagement to represent the associated position may have been boosted for the integration task. This is in line with empirical and computational modeling works that have shown the manipulation of activity-silent representations requires reinstatement into persistent neural activity (Trübutschek et al., 2019), with the strength of this activity scaling with the degree of manipulation required (Masse et al., 2019). The preparatory difference is further compatible with the dual mechanisms of the cognitive control framework (Braver, 2012): integration may have encouraged a proactive strategy, whereas recognition may have afforded a more reactive approach. This proactive process would allow establishing item-rule associations, which have previously been shown to trigger the reactivation of task-related information from LTM (Şentürk et al., 2024). Our results extend this literature principle to the LTM domain, providing direct neural evidence that WM represents LTM content more strongly for mental operations compared to recognition, albeit transiently.

Importantly, despite brief differences, reactivation levels mostly converged across tasks over the retention interval and during operations, suggesting that both integration and recognition ultimately required comparable WM engagement for storing LTM representations. WM engagement for the novel positions was likewise equivalent across tasks. Together, these findings indicate that the task-relevant representations were maintained at comparable strengths for both tasks. Accordingly, heightened WM activity reported in previous studies of mental manipulation may reflect procedural demands of mental operations rather than stronger representations of task-relevant contents.

Having characterized the content-level maintenance before and during the mental operation, we next examined whether the integration and recognition tasks differed in cognitive control related processing, as reflected in frontal theta power. Frontal theta has been associated with the recruitment of cognitive control (Cavanagh & Frank, 2014) and with successful manipulation in WM (Itthipuripat et al., 2013). We found no difference in frontal theta power between the Integration and Recognition tasks throughout the trial, providing no evidence that the tasks differed in cognitive-control demands. Although this may contrast with previous evidence that manipulation places greater demands on control than maintenance (Veltman et al., 2003; D’Esposito et al., 1999), some recent fMRI findings suggest that manipulation recruits distinct networks from those for maintenance rather than uniform increases in overall activity. For example, Mohr et al. (2006) and Eldreth et al. (2006) found differential recruitment of frontal regions across the two operations, with some regions increasing and others decreasing in activity. Similarly, Davis et al. (2018) identified distinct maintenance and manipulation networks whose dissociation increases as their corresponding demands grew within a single task. Thus, equivalent frontal theta power need not imply identical neural processing: spatially distinct or oppositely varying neural contributions may produce no detectable difference in scalp-level theta power. Consistent with this possibility, here, alpha-band MVPA distinguished the integration and recognition tasks based on their distributed activity patterns despite no difference in overall alpha power. Overall, the neural distinction between manipulation vs maintenance may therefore be expressed through differences in the pattern of activity rather than a simple increase in the overall level of activity reflecting cognitive control.

Beyond the degree of reactivation and cognitive demand, we tested whether the representational format of LTM content differed across tasks by training an IEM on one task and testing it on the other. Cross-training showed above-chance CTF slopes, indicating that spatial representations were maintained in an overlapping format across integration and recognition blocks that remained stable regardless of task context. Thus, representational formats of spatial information stored for recognition and mental operations are comparable. This builds on our former work showing dynamically updated spatial WM representations share overlapping formats with non-updated ones (Günseli et al., 2024), and extends it to information stored for mental operations.

One possible limitation is overpractice: with extensive practice on the same LTM positions, the integration operation may have become automatized, potentially accounting for the absence of frontal theta differences and the convergence of reactivation levels. However, the accuracy difference between tasks persisted throughout the experiment, and behavioral costs continued to scale with the magnitude of spatial transformation (Figure 2C&S1), arguing against full automatization.

A second consideration is that the integration task allowed participants to begin the mental operation immediately after the second position was presented, whereas recognition had no comparable need to act on the information until probe onset. This temporal asymmetry could itself produce an early reactivation difference independent of any difference in WM engagement, since integration’s information was immediately task-relevant while recognition’s was not. However, this account does not fully explain our results: although delayed relative to integration, LTM reactivation in the recognition task also emerged well before probe onset, indicating that the relevant positions were reactivated in preparation for both tasks, not only at the moment they were needed for comparison.

Third, the low memory load in our task (two positions) may constrain the degree to which participants rely on latent LTM representations, as WM may not need LTM’s support when load is low (Bartsch et al., 2024), leaving less room for offloading differences to emerge between tasks. However, recent work shows that anticipated memory load being 2 or 4 does not affect the extent to which participants rely on active WM versus latent LTM states (Yılmaz et al., 2026), suggesting that low memory load may not be a likely explanation for the lack of condition differences.

Lastly, our findings are restricted to the spatial domain. Whether the same pattern holds for non-spatial features, such as color or object identity, or for non-spatial operations, remains an open question.

In summary, this study provides neural evidence that WM involvement in mental operations on LTM information is not uniformly elevated relative to recognition. There was no increase in demands for cognitive control as indexed by frontal theta power, and the differences in the strength of task-relevant representations were not sustained throughout the trial. Thus, the need for mental operations may not be a factor that drives stronger WM engagement relative to mere recognition. Overall, these findings contribute to our understanding of how WM is recruited for storing and operating on information retrieved from LTM.

## Author Contributions

Conceptualizing and designing the experiment; İ. Efsane Algın, Eren Günseli. Collecting the experimental data; İ. Efsane Algın. Conducting the analyses; Eren Günseli, İ. Efsane Algın. Writing the paper; İ. Efsane Algın, Eren Günseli. Supervision; Eren Günseli.

## Acknowledgements

We would like to thank our research assistants Yağmur Zehra Yılmaz, Cem Dereli, Ece Lüle, Meral Özkütük, Rime Elkherrat and Mahmut Kaya for their assistance during data collection.

## Citation Diversity Statement

We value diversity in scholarly citations but did not assess the gender or other diversity characteristics of our citations.

## Declarations

### Funding

The Scientific and Technological Research Council of Türkiye (TÜBİTAK) Incentive Award and the Turkish Academy of Sciences Young Investigator Award to E.G., and a TÜBİTAK 2210-A National M.Sc. Scholarship to İ.E.A.

### Conflicts of Interest

The authors declare no competing financial interests.

### Ethics Approval

The study was performed according to the Declaration of Helsinki principles and the ethics approval was granted by the Sabancı University (SUREC) ethics committee.

### Data and Code Accessibility

All data, analysis code, and experimental materials are available at OSF: https://osf.io/2sxqr

---

**Figure S1.**
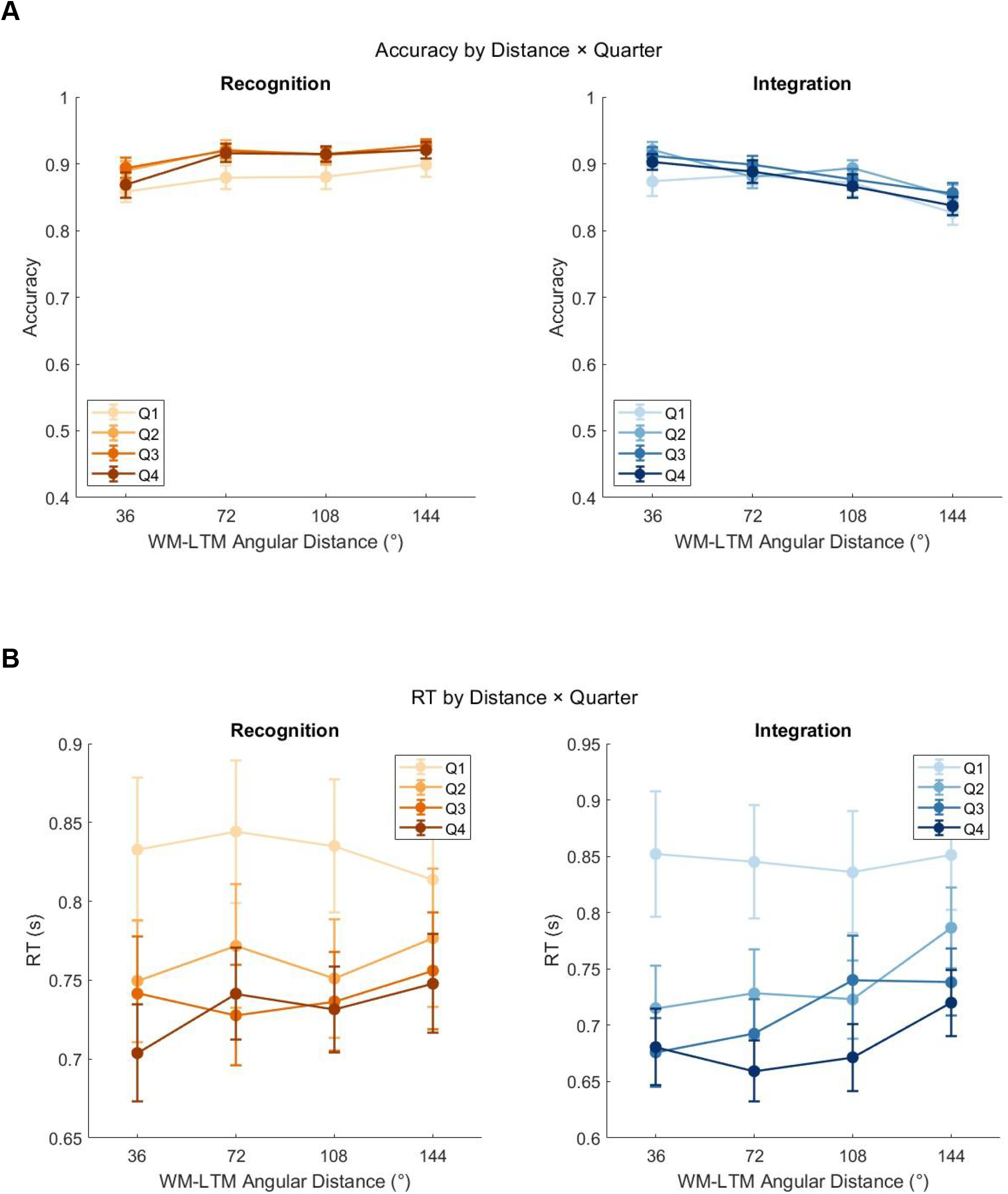
Accuracies and Reaction times by Angular Distance per experimental quarter. Behavioral performance as a function of WM–LTM angular distance, split by experimental quarter (Q1–Q4) and shown separately for the Recognition and Integration tasks. **A.** Accuracy. **B.** Reaction time. Darker shades indicate later quarters (orange shades for Recognition, blue shades for Integration). The effect of angular distance was consistent across quarters in both tasks. Error bars denote ±1 SEM

**Figure S2.**
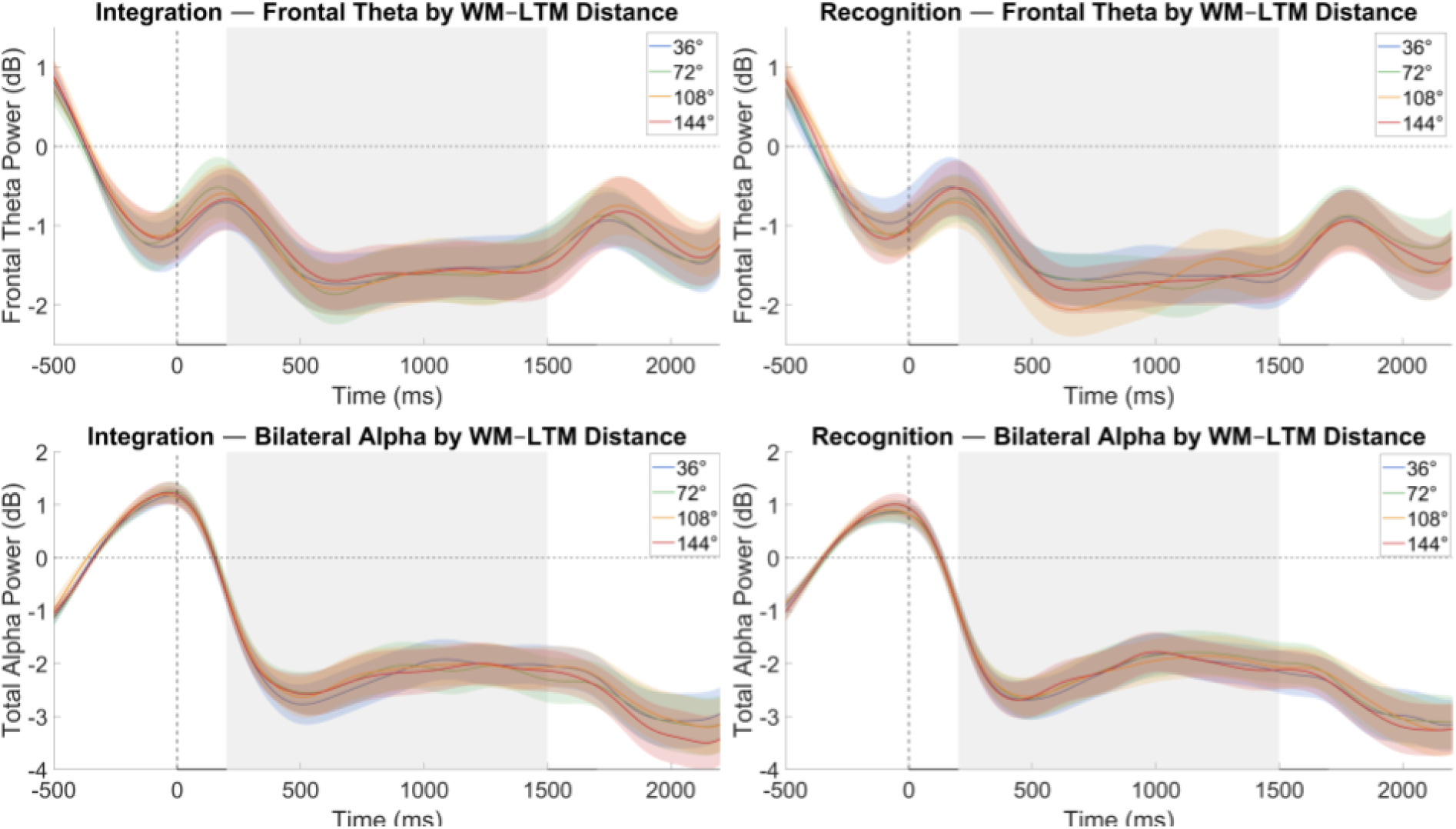
Frontal theta and bilateral alpha power by WM–LTM distance. Frontal theta power (top) and bilateral alpha power (bottom) as a function of WM–LTM angular distance (36°, 72°, 108°, 144°), shown separately for the Integration (left) and Recognition (right) tasks. Bilateral alpha was modulated by angular distance, with no interaction with task type, whereas frontal theta showed no modulation. Shading denotes ±1 SEM; grey marks the retention interval (200–1500 ms).

**Figure S3.**
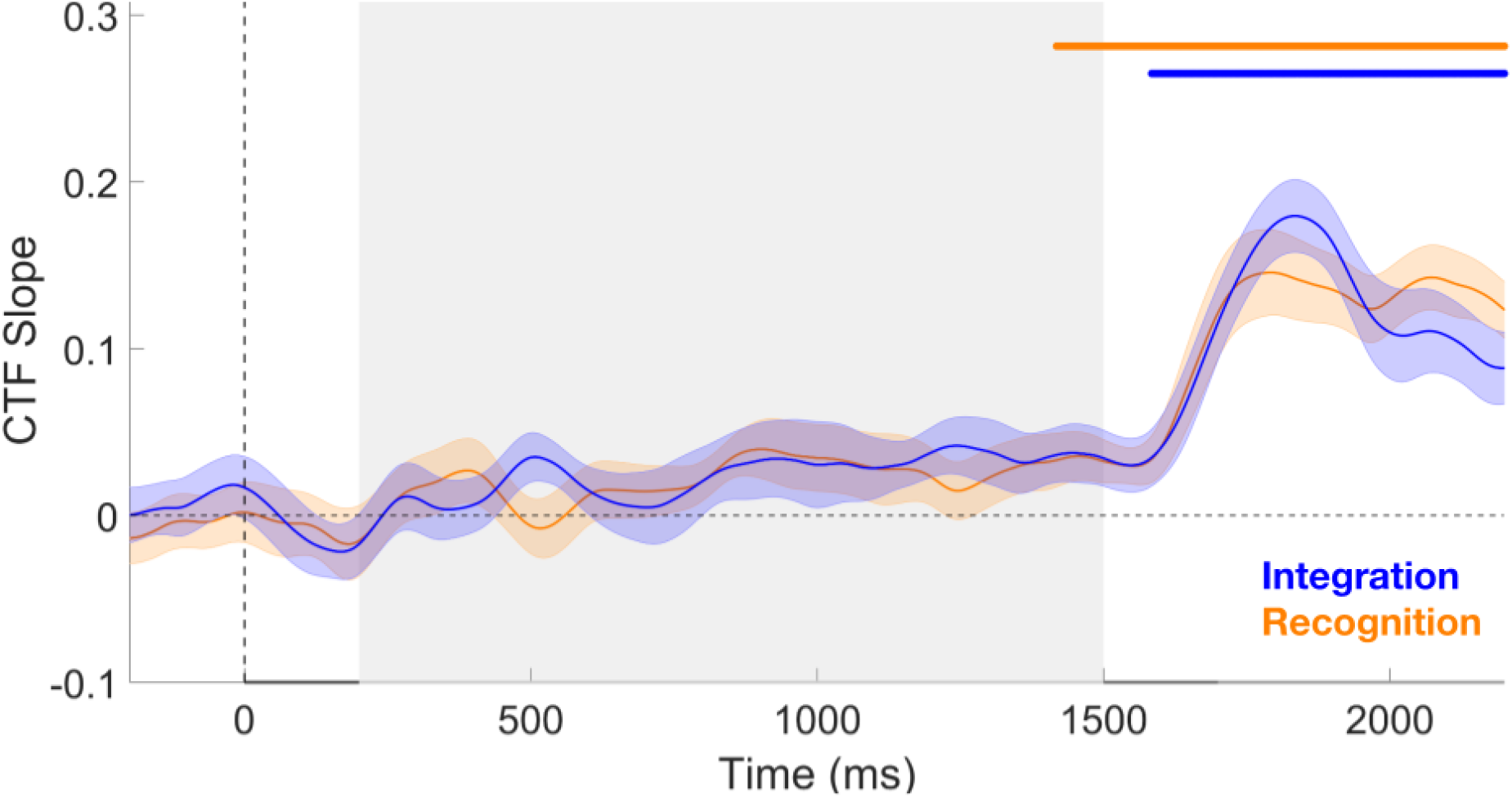
Within-Task Comparisons of Midpoint Position CTF Slopes Against Null. To complement the retention-interval average analysis reported in text S2, we used cluster-based permutation testing to examine the time course of Midpoint Position CTF slopes against permuted (null) CTF slopes within each task condition. For the recognition task, one significant positive cluster emerged from 1500 to 2200 ms (p <.05; two-tailed). For the integration task, one significant positive cluster emerged from 1584 to 2200 ms (p <.05; two-tailed)

**Table S1.** Paired samples test for probe types in the Recognition Task for Accuracy and Reaction Times.

| Accuracy |  |  |  |  |  |
| --- | --- | --- | --- | --- | --- |
| Condition | BF<sub>10</sub> | 95% Credible Interval | *t*(24) | *p* | Cohen's *d* |
| LTM vs WM | 11.81 | [0.174 1.023] | 3.244 | .003* | 0.649 |
| Reaction Times |  |  |  |  |  |
| Condition | BF<sub>10</sub> | 95% Credible Interval | *t*(24) | *p* | Cohen's *d* |
| LTM vs WM | 18.32 | [-1.069 -0.207] | -3.451 | .002* | -0.690 |

## Text S1

In our exploratory behavioral analysis we observed that the computational effort required for the spatial operation scaled with the magnitude of the transformation. To examine whether this behavioral pattern was accompanied by corresponding changes in neural markers, we tested whether frontal theta power and bilateral alpha differed across angular distances (operational difficulty) for both tasks using frequentist and Bayesian Repeated Measures ANOVA with a 2 (Recognition vs Integration) x 4 (36, 72, 108 and 144 degree) design and assessed the effects by comparing across all models (Figure S2).

For bilateral alpha, there was a main effect of angular distance, BF₁₀ = 8.75, *F*(3, 72) = 6.42, *p* <.001, but without an interaction with the task type, BF₀₁ = 7.6, *F*(3, 72) = 0.87, *p* =.46, possibly reflecting attentional engagement to a wider area. Furthermore, Frontal Theta was not modulated by angular distance BF₀₁ = 24.68, *F*(2.49, 59.72) = 0.82, *p* =.47, without an interaction between task types, BF₀₁ = 52.69, *F*(2.01, 48.24) = 1.12, *p* =.34, further supporting the possibility that this neural signature may not be fully sensitive to the computational demands of the operation itself, rather confounded by other factors such as difficulty, or load.

## Text S2

Lastly, we examined whether and how the midpoint positions are represented in WM. Because the 20 possible midpoint positions yielded insufficient trial counts per position for stable IEM estimation, we grouped them into five bins of four adjacent positions. We repeated this grouping with four offset schemes and averaged the resulting CTF slopes. To assess whether midpoint selectivity was specific to the integration task, we also computed the midpoint of the two memory positions for recognition trials and repeated the same IEM procedure. CTF slopes were averaged across the 1700–2200 ms second retention interval.

The resulting CTF slopes are plotted in Figure S3. A Bayesian t-test provided moderate evidence for equivalent midpoint selectivity across the two tasks, BF₀₁ = 4.66, t(24) = 0.2, *p* =.84. Thus, channel responses tuned to the midpoint position were not specific to the integration task but similarly present during recognition. One possibility is that participants automatically computed the midpoint even when the task did not require it. We consider this unlikely given the distinct IEM and MVPA profiles observed for the two task types, as well as the dissociable effects of angular distance on accuracy and RT (see Figure 2, S1). Instead, the equivalent midpoint selectivity may reflect a property of simultaneous spatial maintenance: when two positions are concurrently held in WM, the peak of the reconstructed tuning curve may gravitate toward their midpoint, producing midpoint-tuned channel responses without an explicit integration operation.

## Notes

https://osf.io/2sxqr

### Competing Interest Statement

The authors have declared no competing interest.

### Summary of Updates

In the previous version, the preprint information text was on top of the figures on pages 12, 13, 16 and 17. In this version, we moved the figures lower on the page to leave space for the information text.

## References

ActiCHamp Plus. (2022). Brain Products GmbH.

Allon, A. S., Balaban, H., & Luria, R. (2014). How low can you go? Changing the resolution of novel complex objects in visual working memory according to task demands. Frontiers in Psychology, 5, 265. 10.3389/fpsyg.2014.00265

Ataseven, N., Algın, İ. E., Yücel, D., Güler, B., Todorova, L., Fukuda, K., & Günseli, E. (2026). Memory reactivation levels remain unaffected by anticipated interference. Journal of Cognitive Neuroscience, 1–12.

Baddeley, A. (2020). Working memory. In A. Baddeley, M. W. Eysenck, & M. C. Anderson, Memory (3rd ed., pp. 71–111). Routledge.

Bartsch, L. M., Frischkorn, G. T., & Shepherdson, P. (2024). When load is low, working memory is shielded from long-term memory’s influence. Journal of Cognition, 7(1), 44. 10.5334/joc.368

BrainVision Recorder (1.24.0001). (2022). Brain Products GmbH.

Braver, T. S. (2012). The variable nature of cognitive control: A dual mechanisms framework. Trends in Cognitive Sciences, 16(2), 106–113. 10.1016/j.tics.2011.12.010

Carlisle, N. B., Arita, J. T., Pardo, D., & Woodman, G. F. (2011). Attentional templates in visual working memory. Journal of Neuroscience, 31(25), 9315–9322.

Cavanagh, J. F., & Frank, M. J. (2014). Frontal theta as a mechanism for cognitive control. Trends in Cognitive Sciences, 18(8), 414–421. 10.1016/j.tics.2014.04.012

Chota, S., & Van der Stigchel, S. (2021). Dynamic and flexible transformation and reallocation of visual working memory representations. Visual Cognition, 29(7), 409–415. 10.1080/13506285.2021.1891168

Davis, S. W., Crowell, C. A., Beynel, L., Deng, L., Lakhlani, D., Hilbig, S. A., Lim, W., Nguyen, D., Peterchev, A. V., Luber, B. M., Lisanby, S. H., Appelbaum, L. G., & Cabeza, R. (2018). Complementary topology of maintenance and manipulation brain networks in working memory. Scientific reports, 8(1), 17827. 10.1038/s41598-018-35887-2

De Pisapia, N., Slomski, J. A., & Braver, T. S. (2007). Functional specializations in lateral prefrontal cortex associated with the integration and segregation of information in working memory. Cerebral Cortex, 17(5), 993–1006.

Delorme, A., & Makeig, S. (2004). EEGLAB: An open source toolbox for analysis of single-trial EEG dynamics including independent component analysis. Journal of Neuroscience Methods, 134(1), 9–21. 10.1016/j.jneumeth.2003.10.009

D’Esposito, M., Postle, B. R., Ballard, D., & Lease, J. (1999). Maintenance versus manipulation of information held in working memory: an event-related fMRI study. Brain and cognition, 41(1), 66–86. 10.1006/brcg.1999.1096

Dienes, Z. (2021). How to use and report Bayesian hypothesis tests. *Psychology of Consciousness: Theory*, Research, and Practice, 8(1), 9–26. 10.1037/cns0000258

Eldreth, D. A., Patterson, M. D., Porcelli, A. J., Biswal, B. B., Rebbechi, D., & Rypma, B. (2006). Evidence for multiple manipulation processes in prefrontal cortex. Brain research, 1123(1), 145–156. 10.1016/j.brainres.2006.07.129

Faul, F., Erdfelder, E., Lang, A. G., & Buchner, A. (2007). G* Power 3: A flexible statistical power analysis program for the social, behavioral, and biomedical sciences. Behavior research methods, 39(2), 175–191.

Foster, J. J., Sutterer, D. W., Serences, J. T., & Awh, E. (2016). The topography of alpha-band activity tracks the content of spatial working memory. Journal of Neurophysiology, 115(1), 168–177. 10.1152/jn.00860.2015

Glahn, D. C., Kim, J., Cohen, M. S., Poutanen, V. P., Therman, S., Bava, S.,…Cannon, T. D. (2002). Maintenance and manipulation in spatial working memory: Dissociations in the prefrontal cortex. NeuroImage, 17(1), 201–213.

Günseli, E., Foster, J. J., Sutterer, D. W., Todorova, L., Vogel, E. K., & Awh, E. (2024). Encoded and updated spatial working memories share a common representational format in alpha activity. iScience, 27(2), 108963. 10.1016/j.isci.2024.108963

Günseli, E., Olivers, C. N. L., & Meeter, M. (2014). Effects of search difficulty on the selection, maintenance, and learning of attentional templates. Journal of Cognitive Neuroscience, 26(9), 2042–2054. 10.1162/jocn_a_00600

Günseli, E. (n.d.). Beyond access: A selective reactivation account of working memory (in prep.)

Hyun, J. S., & Luck, S. J. (2007). Visual working memory as the substrate for mental rotation. Psychonomic Bulletin & Review, 14, 154–158. 10.3758/BF03194043

Itthipuripat, S., Wessel, J. R., & Aron, A. R. (2013). Frontal theta is a signature of successful working memory manipulation. Experimental Brain Research, 224(2), 255–262. 10.1007/s00221-012-3305-3

JASP Team. (2019). JASP (Version 0.95.2) [Computer software]. https://jasp-stats.org/

Lee, M. D., & Wagenmakers, E.-J. (2014). Bayesian cognitive modeling: A practical course (1st ed.). Cambridge University Press. 10.1017/CBO9781139087759

Liu, B., Li, X., Theeuwes, J., & Wang, B. (2022). Long-term memory retrieval bypasses working memory. NeuroImage, 261, 119513. 10.1016/j.neuroimage.2022.119513

Logie, R. H. (2003). Spatial and visual working memory: A mental workspace. In Psychology of Learning and Motivation (Vol. 42, pp. 37–78). Academic Press.

Logie, R. H., Gilhooly, K. J., & Wynn, V. (1994). Counting on working memory in arithmetic problem solving. Memory & Cognition, 22(4), 395–410.

Lopez-Calderon, J., & Luck, S. J. (2014). ERPLAB: An open-source toolbox for the analysis of event-related potentials. Frontiers in Human Neuroscience, 8. 10.3389/fnhum.2014.00213

Machizawa, M. G., Goh, C. C., & Driver, J. (2012). Human visual short-term memory precision can be varied at will when the number of retained items is low. Psychological Science, 23(6), 554–559. 10.1177/0956797611431988

Maris, E., & Oostenveld, R. (2007). Nonparametric statistical testing of EEG-and MEG-data. Journal of Neuroscience Methods, 164(1), 177–190. 10.1016/j.jneumeth.2007.03.024

Master, S. L., Li, S., & Curtis, C. E. (2024). Trying harder: How cognitive effort sculpts neural representations during working memory. Journal of Neuroscience, 44(28), e0060242024. 10.1523/JNEUROSCI.0060-24.2024

Masse, N. Y., Yang, G. R., Song, H. F., Wang, X. J., & Freedman, D. J. (2019). Circuit mechanisms for the maintenance and manipulation of information in working memory. Nature Neuroscience, 22(7), 1159–1167.

Mohr, H. M., Goebel, R., & Linden, D. E. (2006). Content-and task-specific dissociations of frontal activity during maintenance and manipulation in visual working memory. Journal of Neuroscience, 26(17), 4465–4471. 10.1523/JNEUROSCI.5232-05.2006

Oostenveld, R., Fries, P., Maris, E., & Schoffelen, J. M. (2011). FieldTrip: Open source software for advanced analysis of MEG, EEG, and invasive electrophysiological data. Computational Intelligence and Neuroscience, 2011, 156869. 10.1155/2011/156869

Oosterhof, N. N., Connolly, A. C., & Haxby, J. V. (2016). CoSMoMVPA: Multi-modal multivariate pattern analysis of neuroimaging data in Matlab/GNU Octave. Frontiers in Neuroinformatics, 10,27. 10.3389/fninf.2016.00027

Risko, E. F., & Gilbert, S. J. (2016). Cognitive offloading. Trends in Cognitive Sciences, 20(9), 676–688. 10.1016/j.tics.2016.07.002

Sauseng, P., Klimesch, W., Doppelmayr, M., Pecherstorfer, T., Freunberger, R., & Hanslmayr, S. (2005). EEG alpha synchronization and functional coupling during top-down processing in a working memory task. Human Brain Mapping, 26(2), 148–155.

Schönbrodt, F. D., & Wagenmakers, E.-J. (2018). Bayes factor design analysis: Planning for compelling evidence. Psychonomic Bulletin & Review, 25(1), 128–142. 10.3758/s13423-017-1230-y

Şentürk, Y. D., Ünver, N., Demircan, C., Egner, T., & Günseli, E. (2024). The reactivation of task rules triggers the reactivation of task-relevant items. Cortex, 171, 465–480. 10.1016/j.cortex.2023.10.024

Sutterer, D. W., Foster, J. J., Serences, J. T., Vogel, E. K., & Awh, E. (2019). Alpha-band oscillations track the retrieval of precise spatial representations from long-term memory. Journal of Neurophysiology, 122(2), 539–551. 10.1152/jn.00268.2019

Trübutschek, D., Marti, S., Ueberschär, H., & Dehaene, S. (2019). Probing the limits of activity-silent non-conscious working memory. Proceedings of the National Academy of Sciences, 116(28), 14358–14367. 10.1073/pnas.1820730116

Unsworth, N., Fukuda, K., Awh, E., & Vogel, E. K. (2014). Working memory and fluid intelligence: Capacity, attention control, and secondary memory retrieval. Cognitive Psychology, 71, 1–26.

Veltman, D. J., Rombouts, S. A., & Dolan, R. J. (2003). Maintenance versus manipulation in verbal working memory revisited: an fMRI study. NeuroImage, 18(2), 247–256. 10.1016/s1053-8119(02)00049-6

Wolff, M. J., Jochim, J., Akyürek, E. G., & Stokes, M. G. (2017). Dynamic hidden states underlying working-memory-guided behavior. Nature Neuroscience, 20(6), 864–871. 10.1038/nn.4546

Yılmaz, Y., Ataseven, N., Kruijne, W., Akyürek, E., & Günseli, E. (2026). Passive but accessible: Studied information is not actively stored in working memory, yet attended regardless of anticipated load. Journal of Cognitive Neuroscience, 1–14.

